# CNView: a visualization and annotation tool for copy number variation from whole-genome sequencing

**DOI:** 10.1101/049536

**Authors:** Ryan L. Collins, Matthew R. Stone, Harrison Brand, Joseph T. Glessner, Michael E. Talkowski

**Affiliations:** Psychiatric and Neurodevelopmental Genetics Unit and Molecular Neurogenetics Unit, Center for Human Genetic Research, Massachusetts General Hospital, Boston, MA 02114, USA; Department of Neurology, Harvard Medical School, Boston, MA 02114, USA; Program in Medical and Population Genetics, Broad Institute, Cambridge, MA 02141, USA

## Abstract

**Summary:** Copy number variation (CNV) is a major component of structural differences between individual genomes. The recent emergence of population-scale whole-genome sequencing (WGS) datasets has enabled genome-wide CNV delineation. However, molecular validation at this scale is impractical, so visualization is an invaluable preliminary screening approach when evaluating CNVs. Standardized tools for visualization of CNVs in large WGS datasets are therefore in wide demand.

**Methods & Results:** To address this demand, we developed a software tool, CNView, for normalized visualization, statistical scoring, and annotation of CNVs from population-scale WGS datasets. CNView surmounts challenges of sequencing depth variability between individual libraries by locally adapting to cohort-wide variance in sequencing uniformity at any locus. Importantly, CNView is broadly extensible to any reference genome assembly and most current WGS data types.

**Availability and Implementation:** CNView is written in R, is supported on OS X, MS Windows, and Linux, and is freely distributed under the MIT license. Source code and documentation are available from https://github.com/RCollins13/CNView

## 1 Introduction

Deletions and duplications of genomic segments, collectively known as copy number variants (CNVs), are the single largest influence in determining the content and organization of an individual genome (Sudamant *et al*., 2015) and are strongly associated with an increased risk of numerous cancers and neurodevelopmental disorders (McCarroll & Altshuler, 2007). Whole genome sequencing (WGS) is the only currently practical method able to capture the full size spectrum of CNV in the human genome (Sudamant *et al*., 2015). Detection of CNV in WGS data commonly relies on measuring relative losses or gains in depth of sequencing coverage, but most algorithms yield too many candidate CNV calls to be molecularly validated at scale. Visual assessment of sequencing depth can quickly assess CNV *in silico*, but there is presently a paucity of tools for CNV visualization from population-scale WGS data.

We present CNView, an R software tool for normalized visualization of sequencing depth in population-scale WGS datasets. CNView applies global intra-sample normalization and localized inter-sample normalization to delineate, annotate, and statistically score CNVs in individual samples or up to hundreds of WGS libraries simultaneously.

## 2 Methods & Application

As input, the BEDtools *coverage* and *uniongbed* commands are used to generate a matrix of uniformly binned sequencing coverages for each library (Quinlan & Hall, 2010; Online Documentation). Compressing coverage into bins of 100bp-1kb smooths visible noise in the sequencing depth while also lowering local computational requirements. After generating this input coverage matrix, CNView can assess and visualize any query region in up to 300 samples at once in under a minute on a laptop with a 2.3 GHz dual-core processor and 8GB RAM.

CNView has six sequential steps: (1) matrix filtering, (2) matrix compression, (3) intra-sample normalization, (4) inter-sample normalization, (5) coverage visualization, and (6) genome annotation. Coverage is extracted from the query region including several flanking megabases (default=5Mb). CNView further compresses this subsetted matrix to reduce local noise. Each library is then normalized by dividing the coverage of each bin by the library’s median nonzero binwise coverage. The intra-sample normalized coverage in each bin is then normalized across all samples to fit the standard normal distribution (μ=0, σ=1). This normalization procedure produces a t-score per sample per bin.

Coverage t-scores are plotted as semi-contiguous step functions for each sample specified by the user. Individual bins significantly depleted or enriched for normalized sequencing depth are indicated by red and blue outlines, respectively (α=0.05, Bonferroni correction). P-values of deletion and duplication are calculated for each highlighted interval by computing the mean t-score of all bins overlapping that interval. The background of each plot is shaded with measurements of central tendency (median) and deviation (median absolute deviation; MAD) per bin. Median and MAD identify regions with unusually high or low coverage variability across samples, which could occur at sites of multiallelic segmental duplications or across regions of heterochromatin, as examples. These features of the coverage distribution per bin cannot be captured by mean and standard deviation due to the normalization function applied in step four, but are readily reflected at regions where the median and MAD diverge significantly from the scaled mean (0) and standard deviation (1). Finally, CNView provides an extensible interface to the UCSC MySQL database and plots specified genomic annotations beneath the normalized coverage signal (Kent *et al*., 2002).

## 3 Results

We previously applied an alpha version of CNView to delineate simple and complex CNVs in two independent WGS cohorts (Brand *et al*., 2014; Brand *et al*., 2015). Here, we also applied CNView to a recently described WGS cohort of 160 individuals comprising 40 quartet families (Turner *et al*., 2016) to show that CNView readily visualizes simple CNVs in individual samples, like the 46kb paternally-inherited, two-exon deletion of *PDE11A* shown in **Figure 1A**. Further, CNView can provide visual confirmation of unbalanced complex genomic rearrangements or compound CNV sites, as shown in **Figure 1B**. In this example, sequencing analysis predicted two large, rare, overlapping CNVs near the p-terminus of chromosome 7: a 467 kb distal deletion and a 449 kb proximal duplication. CNView assessment of this site provides supporting evidence of the compound CNV by illustrating copy loss of the deletion-specific interval, copy gain of the duplication-specific copy number, and no change in copy number in the overlapping interval between the deletion and duplication. Links to the data used to create both panels of **Figure 1** are available in the **Supplementary Information**.

**Figure 1.**
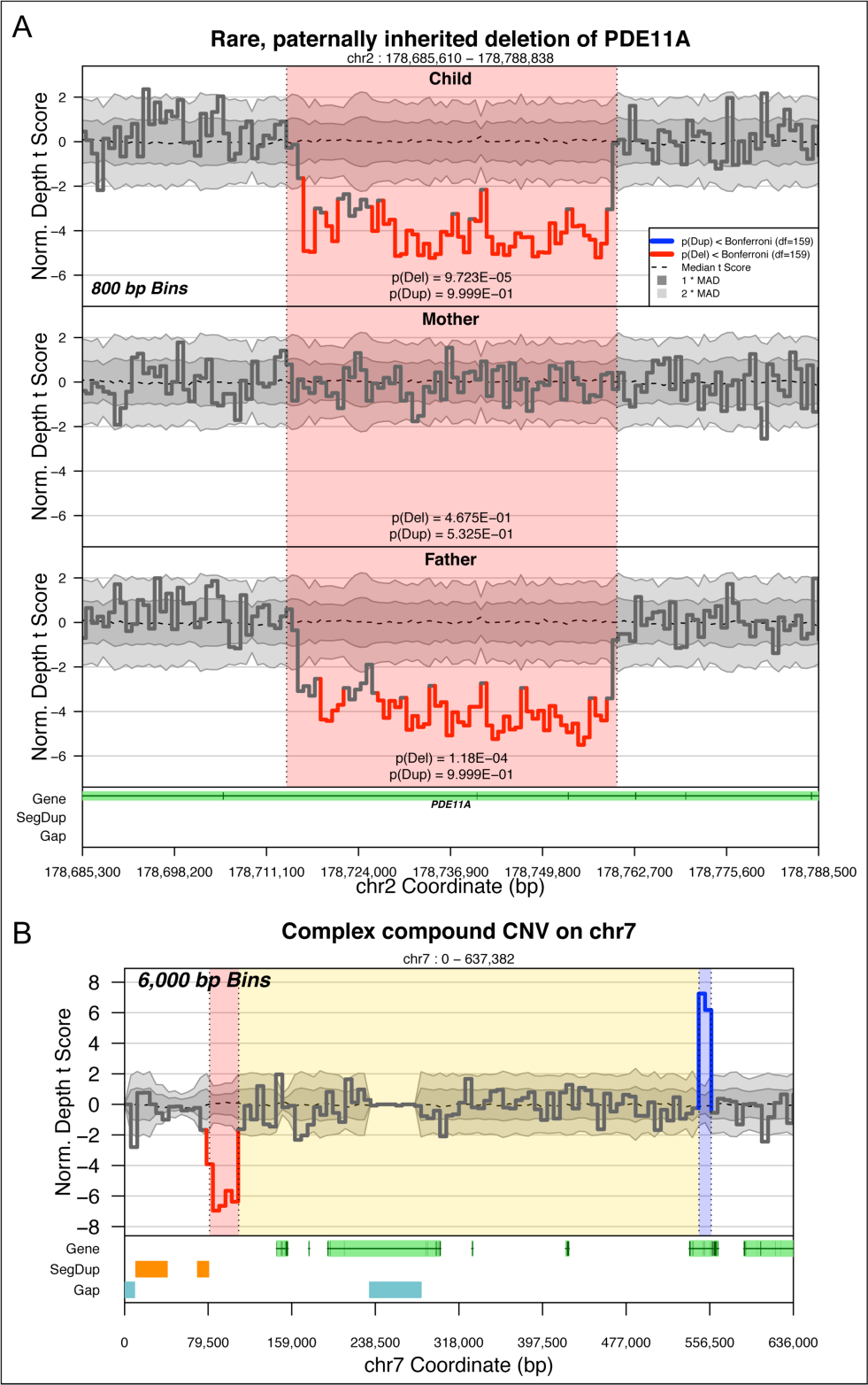
CNView renderings of simple and compound CNVs. Both panels can be regenerated directly from CNView with no post-hoc modifications directly from the **Supplementary Data**. (A) CNView shows copy loss over a 46,166 bp two-exon deletion of *PDE11A* (red) in a child (top) and his father (bottom), but not the mother (middle). (B) CNView provides evidence of a compound CNV, in which a 467 kb deletion overlaps a 449 kb duplication, resulting in small segments of decreased (red) and increased (blue) copy number, while the rest of the site (yellow) remains copy number-neutral (grey).

## Supplementary Information

Supplementary data are available online.

## References

Brand, H. et al. (2014) Cryptic and Complex Chromosomal Abberations in Early-Onset Neuropsychiatric Disorders. Am. J. Hum. Genet., 95(4), 451–461.

Brand, H. et al. (2015) Paired-Duplication Signatures Mark Cryptic Inversions and Other Complex Structural Variation. Am. J. Hum. Genet., 97, 170–176.

Kent, W.J. et al. (2002) The Human Genome Browser at UCSC. Genome Res., 12, 996–1006.

McCarroll, S. and Altschuler, D. (2007) Copy-Number Variation and Association Studies of Human Disease. Nat. Genet., 39, 37–42.

Quinlan, A.R. & Hall, I.M. BEDTools: A Flexible Suite of Utilities for Comparing Genomic Features. Bioinformatics, 26, 841–842.

Sudmant, P. et al. (2015) An Integrated Map of Structural Variation in 2,504 Human Genomes. Nature, 526, 75–81.

Turner, T.N. et al. (2016) Genome Sequencing of Autism-Affected Families Reveals Disruption of Putative Noncoding Regulatory DNA. Am. J. Hum. Genet., 98, 58–74.

